# HIPSTR: highest independent posterior subtree reconstruction in TreeAnnotator X

**DOI:** 10.1101/2024.12.08.627395

**Authors:** Guy Baele, Luiz M. Carvalho, Marius Brusselmans, Gytis Dudas, Xiang Ji, John T. McCrone, Philippe Lemey, Marc A. Suchard, Andrew Rambaut

## Abstract

In Bayesian phylogenetic and phylodynamic studies it is common to summarise the posterior distribution of trees with a time-calibrated consensus phylogeny. While the maximum clade credibility (MCC) tree is often used for this purpose, we here show that a novel consensus tree method – the highest independent posterior subtree reconstruction, or HIPSTR – contains consistently higher supported clades over MCC. We also provide faster computational routines for estimating both consensus trees in an updated version of TreeAnnotator X, an open-source software program that summarizes the information from a sample of trees and returns many helpful statistics such as individual clade credibilities contained in the consensus tree. HIPSTR and MCC reconstructions on two Ebola virus and two SARS-CoV-2 data sets show that HIPSTR yields consensus trees that consistently contain clades with higher support compared to MCC trees. The MCC trees regularly fail to include several clades with very high posterior probability (≥ 0.95) as well as a large number of clades with moderate to high posterior probability (≥ 0.50), whereas HIPSTR achieves near-perfect performance in this respect. HIPSTR also exhibits favorable computational performance over MCC in TreeAnnotator X. Comparison to the recently developed CCD0-MAP algorithm yielded mixed results, and requires more in-depth exploration in follow-up studies. TreeAnnotator X – which is part of the BEAST X (v10.5.0) software package – is available at https://github.com/beast-dev/beast-mcmc/releases.

## 1 Introduction

Bayesian phylogenetic inference remains one of the most widely used frameworks to estimate the evolutionary relationship between a set of genetic or genomic sequences [Larget and Simon, 1999, Huelsenbeck et al., 2001, Ronquist et al., 2012, Suchard et al., 2018, Bouckaert et al., 2019, Höhna et al., 2016]. One of the key outcomes of such a Bayesian analysis is a set of phylogenetic trees sampled from the model posterior. This set is then summarized into an easily disseminable and interpretable result – usually a single representative tree used as a framework to display important phylogenetic relationships and other quantities of interest such as divergence times or trait evolution. Given that this set of trees may contain (tens of) thousands of unique topologies, a range of summary tree calculation methods have been developed over the past decades. An extensive review of these methods is beyond the scope of this paper, but can be found in – for example – Bryant et al. [2003].

Current Bayesian phylogenetic software packages undertake a random walk through the space of trees, usually employing Metropolis-Hastings sampling [Metropolis et al., 1953, Hastings, 1970] to attempt to sample trees from the posterior distribution as this space is explored. Were time and electricity unlimited, a preferred point-estimate of the phylogenetic tree would be the one most frequently visited (the maximum *a posteriori* or MAP tree). In practice, for all but trivially small data sets, these stochastic algorithms will likely never visit the same tree twice. Consequently, the approach taken is to consider constituent parts of the tree independently, reporting the frequency of individual clades (for rooted trees) or splits (for unrooted trees). Hereon we will refer to clades as BEAST [Suchard et al., 2018, Bouckaert et al., 2019] is exclusively focused on rooted trees. For many phylogenetic questions, these clade frequencies can be used directly to provide support for competing hypotheses without considering the tree as a whole. Similarly, estimates of parameters of interest in the models employed (e.g., rates of evolution, substitution model parameters or population size dynamics) are marginalized or averaged over all sampled trees.

However, in many cases it is desirable to represent the totality of the phylogenetic information in the form of a single tree, ‘annotated’ with individual clade frequences and averages or credible intervals of continuous parameters of the tree such as node ages. Furthermore, this tree can be used to visualize jointly-estimated results such as trait evolution or spatial spread. As such, it is essential that this ‘summary’ tree includes all of the highly-supported clades.

The traditional approach to constructing a summary tree, one that long precedes the rise of Bayesian approaches, is the majority-rule consensus tree [Margush and McMorris, 1981]. Often employed to summarize resampling approaches such as bootstrapping [Felsenstein, 1985] or jacknifing [Farris et al., 1996] with maximum-likelihood or maximum-parsimony phylogenetics, this is a tree constructed from a set of clades and their frequencies. The most popular version is the 50% majority consensus tree, a tree constructed such that it contains all of the clades with at least 50% frequency (a strict consensus tree contains only 100% frequency clades). However, for non-trivial data sets these trees will not be fully resolved (bifurcating) as clades that do not meet the criteria for inclusion are collapsed into polytomies. As a result, these methods generally preclude appropriate presentation of time scales or reconstruction of geographic dispersal.

To address these limitations for analyses, where the phylogeny itself is not exclusively the result-of-interest, the BEAST packages [Suchard et al., 2018, Bouckaert et al., 2019] took the approach of finding the maximum clade-credibility (MCC) tree to use as the single tree representative of those sampled. The MCC tree is that amongst the sampled set which has the highest product of all the individual clade frequencies. Thus, it is a tree that the Markov chain actually visited although, in practice, only a small sample of trees is evaluated. For example, by default, BEAST stores a sample of 10 000 trees regardless of the length of the chain. This ‘thinning’ or downsampling is done to remove the autocorrelation that exists between adjacent samples and to produce a tractable set of trees in terms of both storage and the feasibility of downstream analyses. A sample of this size will likely capture all high-frequency clades but will not resolve the relative support for low-frequency clades. Furthermore, the MCC tree may be missing some clades that have less than 100% frequency if, by chance, they don’t all co-occur in at least one tree of this limited sample. Sampling more frequently will not necessarily abrogate this issue because this larger set of trees will have greater autocorrelation.

We here propose a summary tree approach – the highest independent posterior subtree reconstruction (HIPSTR) algorithm – that attempts to address the limitations of both majority-rule consensus trees and MCC trees. HIPSTR aims to construct a tree that contains all the highest frequency, mutually compatible, clades even if that specific tree was never actually sampled by the MCMC.

Since implementing the approach described here, related work has been presented by Berling et al. [2024] which is based on the conditional clade distribution (CCD) which offers an advanced estimate of the posterior probability distribution of the tree space. The authors extend the applicability of CCDs by introducing a new parametrization for CCDs and describing fixed-parameter tractable algorithms to compute the tree with highest probability. Of specific interest to the method we present here is the CCD0-MAP consensus tree, which Berling et al. [2024] recommend as the preferred point estimator for Bayesian phylogenetic inference of time trees. Comparison of HIPSTR to CCD0-MAP is however inhibited by numerical and computational issues that prevent the computation of the CCD0-MAP tree for the large data sets we consider – for these we are compelled to resort to the CCD1-MAP algorithm (see Berling et al. [2024]).

## 2 Methods

### 2.1 Algorithm

An initial pass of the full set of trees collates a table of all observed clades, their frequency (clade credibility) and a list of all observed pairs of child clades. Starting at the root clade of all tips a post-order traversal is performed of this tree structure down to the individual tips. On the return of the traversal a credibility score is computed for each subtree explored. This subtree credibility is the maximum product of the credibility of each pair of descendant subtrees and the credibility of the clade that contains them. As these individual subtrees will be present in many places in the data structure, the maximum subtree credibility and the associated pair of subtrees is stored in a cached keyed by a clade-specific hash for rapid recall. Finally a second post-order traversal of just the maximum credibility subtrees is performed to construct the single, fully bifurcating, HIPSTR tree. A second pass of the set of trees can then accumulate distributions of parameters such as node ages, evolutionary rates, and trait values for the set of clades present in the HIPSTR tree.

### 2.2 Data

We assess the performance of HIPSTR on four complete-genome data sets: two Ebola virus (EBOV) data sets, one containing 1 610 genomes from the 2013-2016 West African EBOV epidemic [Dudas et al., 2017], and another containing 516 genomes from the 2018-2020 Nord Kivu EBOV epidemic [Kinganda-Lusamaki et al., 2021]; and two SARS-CoV-2 data sets, one containing 3 959 genomes from across Europe [Lemey et al., 2021], and another containing 15 616 genomes from the United Kingdom [du Plessis et al., 2021]. We selected these data sets because of their different dimensions (see Table 1) and because they represent key pathogens that continue to pose significant threats to public health. We performed visualisations in baltic v.0.3.0 (https://github.com/evogytis/baltic).

**Table 1:**
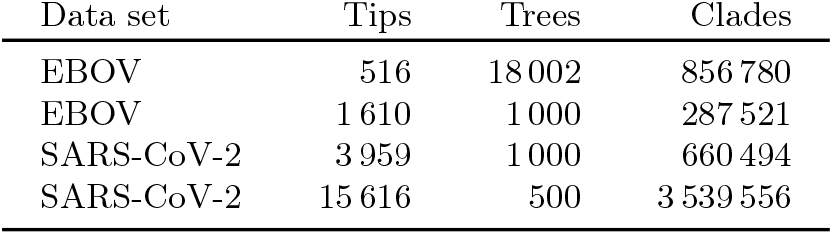
Genomic data set properties and posterior tree sample properties of the four data sets analysed in this study.

Further, based on our findings (but see the Results section) and our extensive experience with the 2013-2016 West African EBOV data set [Dudas et al., 2017], we created a simulated EBOV data set (EBOV-Sim) based on the HIPSTR tree of the original EBOV Bayesian phylogenetic analysis (https://github.com/ebov/space-time/tree/master/Analyses/Phylogenetic). We simulated a sequence data set of 20 000 bp under an HKY substitution model with among-site rate heterogeneity [Hasegawa et al., 1985, Yang, 1996] and a strict clock with an evolutionary rate of 1E-3 substitutions per site per year. We then inferred 10 000 posterior trees from a 100 million iterations analysis in BEAST 1.10.4 [Suchard et al., 2018] under a constant population size model, using the same models as were used to generate the simulated data set.

### 2.3 Computational aspects

The initial implementation of HIPSTR dates back to September 2019 (https://github.com/beast-dev/beast-mcmc/commit/1d3df0eabe2bd133617cf48e9e05eaa810c88152) and was originally called MMCC for maximum marginal clade credibility. Since then, we have restructured important parts of the TreeAnnotator code (part of the upcoming release of BEAST X - v10.5.0) in a more modular form and reimplemented parts of the calculations, aimed at especially benefiting consensus tree construction performance for large file sizes. This implementation benefits the computational performance of both MCC and HIPSTR over the previous version of TreeAnnotator in BEAST 1.10.4 [Suchard et al., 2018] (data not shown).

MCC and HIPSTR calculations were performed in TreeAnnotator X (v10.5.0) using the default run-time settings, whereas the CCD0-MAP and CCD1-MAP calculations in TreeAnnotator v2.7.7 required the allocation of an initial Java heap size of 32 Gb and a maximum Java heap size of 64 Gb. All calculations were performed on an Apple M2 Ultra 24-core processor with 192 Gb of memory.

## 3 Results

In Table 2, we show the performance comparison of HIPSTR over MCC in terms of consensus tree construction and computational cost on the four examples. Of note, phylogenetic analyses with over a thousand genomes resort to storing fewer posterior tree samples, owing to markedly increasing file sizes and ensuing post-processing issues. HIPSTR consistently yields consensus trees with higher log marginal clade credibility and mean individual clade credibility over MCC trees, while doing so in a markedly shorter time compared to the MCC algorithm. Further, HIPSTR consistently includes highly supported clades, while MCC regularly misses out on clades with ≥ 0.95 and occasionally even ≥ 0.99 posterior probability. On the data sets tested HIPSTR performs up to 2× faster than MCC. With increasing numbers of genomes – millions in the case of SARS-CoV-2 – being used in phylogenetic inference, computational performance for consensus tree methods needs to be considered.

**Table 2:** Consensus tree reconstruction and computational performance of MCC, HIPSTR, CCD0-MAP and CCD1-MAP on four genomic data sets (number of sequences between brackets). HIPSTR consistently yields consensus trees with higher log marginal clade credibility, median individual clade credibility and number of clades with ≥50% posterior probability included over MCC, and equal or higher number of clades with ≥99% and ≥95% posterior probability. HIPSTR is also computationally more efficient than MCC, yielding up to 2× higher performance (s: seconds; h: hours). CCD0-MAP could only be computed successfully on our smallest data set but yielded the best performance, at the expense of a very lengthy run time; CCD1-MAP performance was far inferior to all other methods tested. log(MCC): log marginal clade credibility; med(ICC): median individual clade credibility. NA: result not available due to TreeAnnotator v2.7.7 producing a numerical error or becoming unresponsive.

| Data set | Method | med(ICC) | ICC ≥ 0.99 | ICC ≥ 0.95 | ICC ≥ 0.50 | log(MCC) | Time |
| --- | --- | --- | --- | --- | --- | --- | --- |
| EBOV<br>(516) | MCC | 0.1555 | 103/103 | 116/116 | 160/177 | -1355.52 | 116s |
|  | **CCD0-MAP** | **0.2784** | 103/103 | 116/116 | **177/177** | **-923.66** | >5h |
|  | CCD1-MAP | 0.1949 | 100/103 | 112/116 | 165/177 | -1752.89 | 235s |
|  | HIPSTR | 0.2751 | 103/103 | 116/116 | 176/177 | -950.08 | **57s** |
| EBOV<br>(1610) | MCC | 0.1780 | 380/381 | 419/421 | 587/634 | -3960.10 | 10s |
|  | CCD0-MAP | NA | NA | NA | NA | NA | NA |
|  | CCD1-MAP | 0.1660 | 368/381 | 404/421 | 568/634 | -4686.60 | 35s |
|  | **HIPSTR** | **0.3120** | **381/381** | **421/421** | **633/634** | **-3039.60** | **6s** |
| SARS-CoV-2<br>(3959) | MCC | 0.0730 | 852/852 | 955/959 | 1214/1325 | -10146.85 | 40s |
|  | CCD0-MAP | NA | NA | NA | NA | NA | NA |
|  | CCD1-MAP | 0.0720 | 846/852 | 948/959 | 1261/1325 | -11540.45 | 107s |
|  | **HIPSTR** | **0.1880** | 852/852 | **959/959** | **1305/1325** | **-8630.85** | **21s** |
| SARS-CoV-2<br>(15616) | MCC | 0.0120 | 2929/2929 | 2992/2995 | 3693/3980 | -57546.68 | 85s |
|  | CCD0-MAP | NA | NA | NA | NA | NA | NA |
|  | CCD1-MAP | 0.0060 | 2929/2929 | 2994/2995 | 3886/3980 | -58636.38 | 369s |
|  | **HIPSTR** | **0.0440** | 2929/2929 | **2995/2995** | **3980/3980** | **-51352.61** | **50s** |
| EBOV-Sim<br>(1610) | True tree | 0.5239 | 620/620 | 658/658 | 819/819 | -1762.67 | 79s |
|  | MCC | 0.4728 | 619/620 | 657/658 | 784/819 | -2610.98 | 150s |
|  | CCD0-MAP | NA | NA | NA | NA | NA | NA |
|  | CCD1-MAP | 0.4726 | 599/620 | 631/658 | 778/819 | -3543.36 | 311s |
|  | **HIPSTR** | **0.5239** | **620/620** | **658/658** | **819/819** | **-1786.64** | **81s** |

Figure 1 shows a tanglegram comparing the MCC and HIPSTR consensus trees for the 516-genome EBOV data set of Kinganda-Lusamaki et al. [2021]. We observe high similarity between both trees, especially in their backbones, due to the high posterior probability (*>* 0.80) that ensures that they become part of both trees. Away from the backbone, we also observe a large number of differences. In order to not clutter Figure 1 with a large number of posterior probability / clade credibility values (of which summary statistics are readily available from TreeAnnotator X), we compare these values for the MCC and HIPSTR trees in more detail in Figure 2, for all four data sets. Figure 2 shows increased divergence between MCC and HIPSTR trees as posterior support wanes, indicative of the MCC trees not including a number of relatively well-supported nodes (0.50 *<* support *<* 0.80) and the HIPSTR trees consistently selecting better supported nodes among those with lower posterior probability (*<* 0.50). Note that a slight trend difference can be observed for the largest SARS-CoV-2 data set for clade credibility values between 0.80 and 1.0, likely owing to a difference in Bayesian inference methodology due to the much increased number of taxa [du Plessis et al., 2021].

**Fig. 1:**
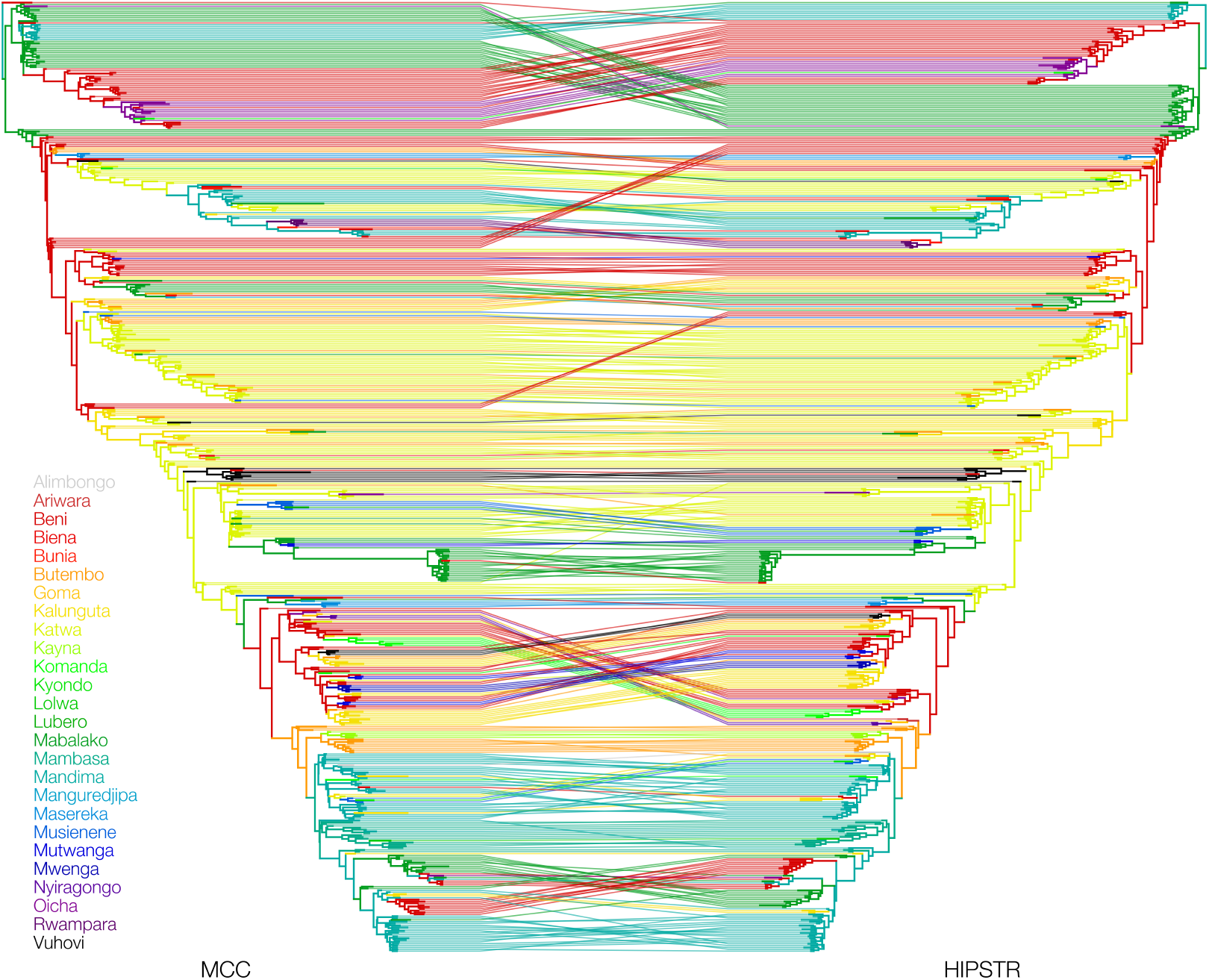
Tanglegram illustrating the similarities and differences between the MCC and HIPSTR trees for a phylogenetic analysis of the 2018-2020 Nord Kivu EBOV data set with 516 complete genomes. The backbone of both trees is highly similar, owing to their high posterior probability (*>* 80), while many differences occur between clusters with lower posterior probability (*<* 50; but see Figure 2). The log product of clade credibilities for the HIPSTR tree (−950.08) is much higher than that of the MCC tree (−1355.52), and only marginally lower than that of the CCD0-MAP tree (−923.66).

**Fig. 2:**
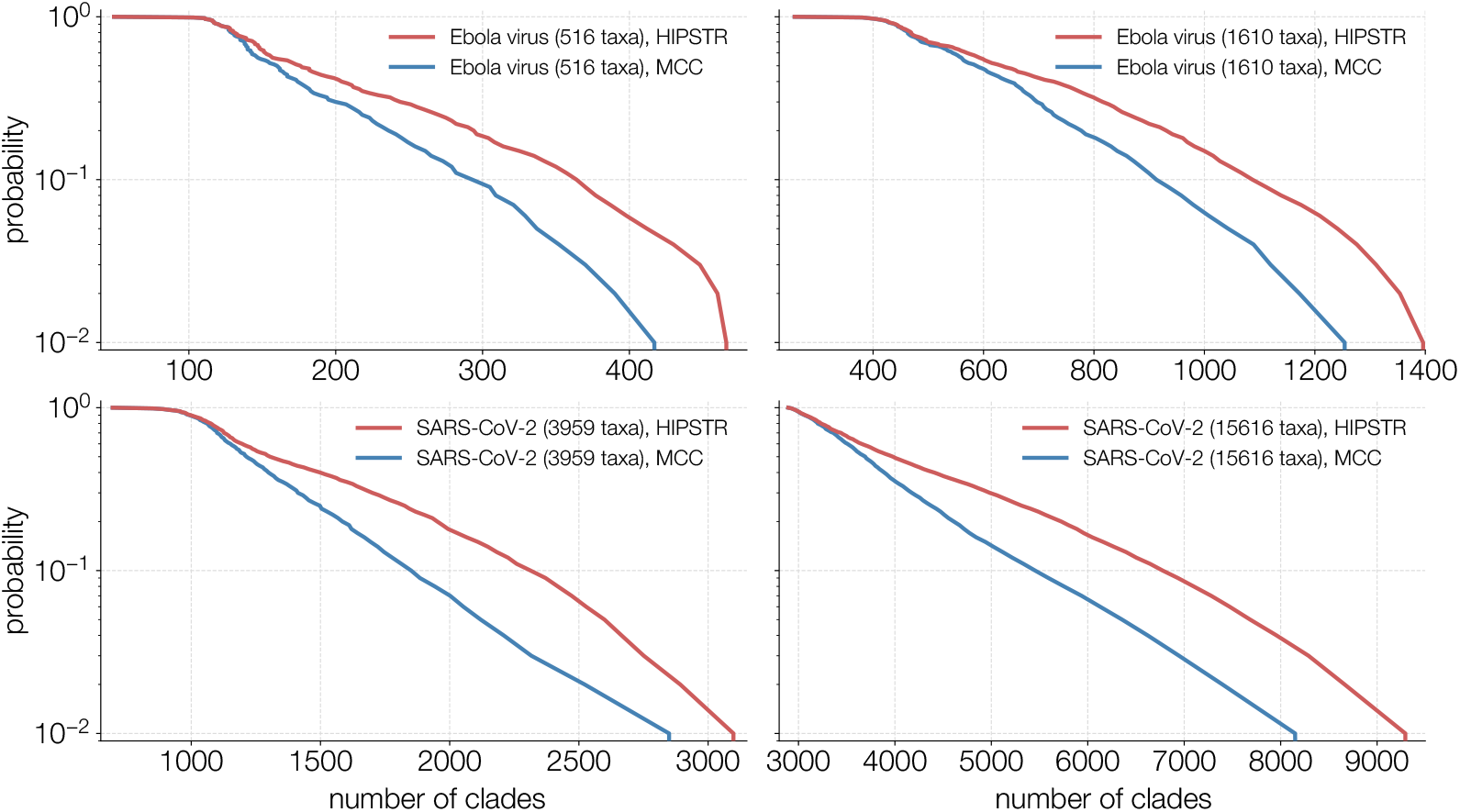
Ordered plots of the posterior probability (log scale) of all clades in the HIPSTR (red) and MCC (blue) trees. Both consensus trees have all the same clades down to approximately 0.80 posterior probability, after which the HIPSTR trees consistently include higher supported clades than the MCC trees – also evident from the differences in log marginal clade credibility; see Table 2.

Comparison of the CCD0-MAP consensus tree [Berling et al., 2024] to the MCC and HIPSTR trees only proved possible for our smallest use case, i.e. the 516-genome EBOV data set [Kinganda-Lusamaki et al., 2021]. Figure 3 shows a tanglegram comparing the HIPSTR and CCD0-MAP consensus trees for this data set, illustrating small differences between the two and only in lower-level clades. As shown in Table 2, only a single clade with posterior probability ≥ 0.50 differs between these two consensus trees, with the CCD0-MAP tree containing this clade. For all other data sets - including our simulated data set - we hence had to resort to the CCD1-MAP consensus tree [Berling et al., 2024] as an alternative. However, in terms of both the log marginal clade credibility and the proportion of clades with posterior probability ≥ 50% this tree is lacking in performance compared to all other consensus tree approaches discussed in this study.

**Fig. 3:**
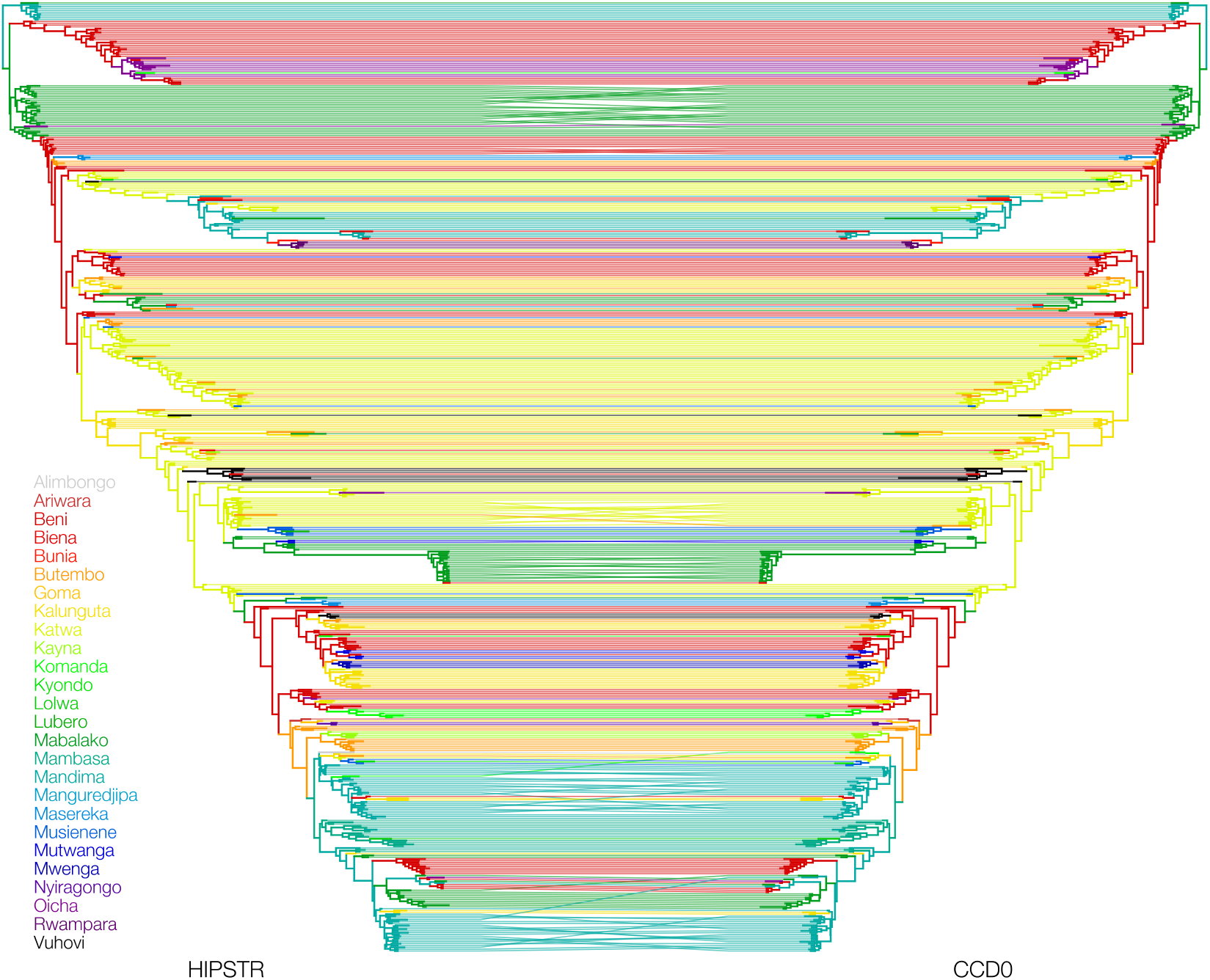
Tanglegram illustrating the similarities and differences between the HIPSTR and CCD0-MAP consensus trees for a phylogenetic analysis of the 2018-2020 Nord Kivu EBOV data set with 516 complete genomes. These highly similar consensus trees have identical backbones and mostly small differences occurring in the lower-level clades. The log product of clade credibilities for the CCD0-MAP tree (−923.66) is slightly higher than that of the HIPSTR tree (−950.08), whereas both are much higher than that of the MCC tree (−1355.52).

## 4 Discussion

We have presented here the novel highest independent posterior subtree reconstruction (HIPSTR) algorithm for reconstructing a potentially unsampled consensus tree from a posterior set of phylogenetic trees, as well as an updated version (X or v10.5.0) of TreeAnnotator. We have shown that HIPSTR consensus trees contain consistently higher clade credibilities than MCC trees, on two EBOV and two SARS-CoV-2 data sets, and that computational performance of HIPSTR surpasses that of MCC reconstruction. Based on these improvements, we recommend the use of the HIPSTR consensus tree over that of the MCC tree.

Comparison of HIPSTR with the CCD0-MAP (employing a parametrization of a CCD based on observed clades) consensus tree [Berling et al., 2024] was only possible for our smallest data set (the 516 genome Ebola virus alignment) due to larger data sets being computationally intractible to the current implementation of the latter method. For the data where CCD0-MAP was successfully constructed, it yielded a further improvement over the HIPSTR tree, including the one clade with ≥ 0.50 posterior probability that the HIPSTR tree did not contain. However, CCD0-MAP calculation came at aconsiderable computational cost of over 5 hours, compared to under 1 minute for the HIPSTR approach. For all other data sets, CCD0-MAP did not produce a result despite signifcant run time and memory being allocated. As an alternative, we calculated the CCD1-MAP (employing a parametrization of a CCD based on observed clade splits and thus having more parameters and requiring more uncorrelated samples) consensus tree for those data sets, but these did not prove competitive with the method proposed here. HIPSTR provides a fast and accurate estimation of the MCC tree that would be obtained from a much larger independent sampling of posterior trees. Hence we conclude that the HIPSTR algorithm and accompanying implementation make for the most appealing choice among available time-calibrated consensus phylogenetic tree approaches and should be preferred over the MCC tree.

## 5 Competing interests

None declared.

## 6 Acknowledgments

G.B. acknowledges support from the Research Foundation - Flanders (“Fonds voor Wetenschappelijk Onderzoek - Vlaanderen,” G0E1420N, G098321N) and from the European Union Horizon 2023 RIA project LEAPS (grant agreement no. 101094685). LMC was partly funded by FAPERJ - Fundação Carlos Chagas Filho de Amparo a Pesquisa do Estado do Rio de Janeiro, Processo SEI 260003/005679/2023 and SEI 260003/013252/2024. G.D. acknowledges the support of European Molecular Biology Organization (EMBO) installation grant IG-5305-2023. X.J. acknowledges support from Louisiana Board of Regents Research Competitiveness Subprogram and the National Science Foundations (NSF) (DEB1754142). P.L., M.A.S. and A.R. acknowledge support from the Wellcome Trust (Collaborators Award 206298/Z/17/Z, ARTIC network), the European Research Council (grant agreement no. 725422 – ReservoirDOCS) and the National Institutes of Health (NIH) (R01 AI153044). M.A.S. acknowledges further support from the NIH through R01 AI162611. P.L. acknowledges support from the Research Foundation, Flanders (“Fonds voor Wetenschappelijk Onderzoek - Vlaanderen,” G066215N, G0D5117N and G0B9317N) and from the European Union Horizon 2020 project MOOD (grant agreement no. 874850). M.B. and G.B. acknowledge support from the DURABLE EU4Health project 02/2023-01/2027, which is co-funded by the European Union (call EU4H-2021-PJ4) under Grant Agreement No. 101102733. Views and opinions expressed are however those of the author(s) only and do not necessarily reflect those of the European Union or the European Health and Digital Executive Agency. Neither the European Union nor the granting authority can be held responsible for them.

## Notes

### Competing Interest Statement

The authors have declared no competing interest.

